# Identification of important amino acid replacements in the 2013-2016 Ebola virus outbreak

**DOI:** 10.1101/075168

**Authors:** Abayomi S Olabode, Derek Gatherer, Xiaowei Jiang, David Matthews, Julian A Hiscox, Stephan Gunther, Miles W Carroll, Simon C Lovell, David L Robertson

**Author notes:** Shared first and. shared last authors (Simon C Lovell, David L Robertson).

## Abstract

The phylogenetic relationships of Zaire ebolavirus have been intensively analysed over the course of the 2013-2016 outbreak. However, there has been limited consideration of the functional impact of this variation. Here we describe an analysis of the available sequence data in the context of protein structure and phylogenetic history. Amino acid replacements are rare and predicted to have minor effects on protein stability. Synonymous mutations greatly outnumber nonsynonymous mutations, and most of the latter fall into unstructured intrinsically disordered regions, indicating that purifying selection is the dominant mode of selective pressure. However, one replacement, occurring early in the outbreak in Gueckedou in Guinea on 31st March 2014 (alanine to valine at position 82 in the GP protein), is close to the site where the virus binds to the host receptor NPC1 and is located in the phylogenetic tree at the origin of the major B lineage of the outbreak. The functional and evolutionary evidence indicates this A82V change likely has consequences for EBOV's host specificity and hence adaptation to humans.

---

The Ebola epidemic of 2013-2016 (named EBOV-Makona) was unprecedented in comparison to previous Ebola virus disease (EVD) outbreaks since 1976, with official estimates at ~29,000 infections and over 11,000 fatalities. The scale of this outbreak is many times larger than any of the previously recorded outbreaks, none of which involved more than 100s of infections. Exploiting contemporary sequencing technologies, researchers have tracked genetic changes via extensive sampling in the affected regions of west Africa^1–8^, generating genome data of EBOV-Makona from hundreds of patient samples. Our evolutionary analysis of the first available dataset^1^ indicated that, despite nucleotide variation, EBOV-Makona is functionally very similar to the variants responsible for previous outbreaks since 1976^9^. Here we extend this analysis to data sets sampled across the outbreak. The unprecedented dimensions of the EBOV-Makona outbreak mean that it is important to monitor for potential signs of viral adaptation to humans.

The evolutionary rates of EBOV in the current West African outbreak have been the subject of discussion^1,3,4,6^ with differences between studies largely due to sampling effects^4^. To understand how EBOV is changing, it is important to distinguish between the underlying mutation rate of the virus (random errors generated during replication) and its substitution rate (mutations that persist after being subject to genetic drift and/or selection). The mutations contributing to the elevated evolutionary rate over shorter timescales^1^ are due to a) incomplete purifying selection and b) neutral nonsynonymous changes that are probably transient and so do not contribute to the substitution rate over longer timescales. With regard to the capacity of EBOV to become adapted to the human population (i.e., endemic) the rate of evolutionary change is less important than the nature of these changes.

EBOV sequence variation must therefore be considered in the context of protein structure and protein-protein interactions. Protein function is tightly linked to protein structure and both are intrinsically linked to protein stability^10,11^ with proteins remaining folded and functional only within a relatively narrow range of temperatures. Non-intuitively, an attenuated EBOV could result in lower viral loads and consequently greater host survival, as long as transmissibility is maintained. For example, the DNA virus myxoma exhibits rapid decreases in virulence due to deleterious mutations that maintained relative transmissibility^12^. Interestingly, there is evidence that EBOV-Makona has lower virulence relative to the 1976 outbreak EBOV-Mayinga^13^; EBOV-Makona viral load is directly linked to survival^14^ and at the population level patient viral loads (and hence virulence) decreased significantly from July to November in 2014^15^. This change may have contributed to increased transmission over the course of the 2013-2016 outbreak. The decrease in virulence might be due to host factors such as genetic differences or alterations in the immune system, or, alternatively, it may result from genetic change in the virus. An attenuated EBOV could be one where amino acid replacements thermodynamically destabilize the protein structure, allowing, for example, high fever in the host to partially control it. Alternatively changes in functional sites could alter the dynamics of infection, for example, as a consequence of more (or less) efficient receptor binding.

To investigate possible EBOV adaptation to humans we have characterised the functional consequences of amino acid residue changes in a structural and evolutionary context across the 2013-2016 outbreak. We find, although relatively rare, both stabilising and destabilising amino acid residue replacements and a tendency for residue changes in the intrinsically disordered regions of EBOV’s proteins. We also find a single variant proximal to the site where the virus binds the host receptor that occurred in Guinea early in the evolution of the outbreak. We discuss the implications of these findings.

## Results

We analysed all complete genome sequence data from the EBOV-Makona outbreak, sampled between March 2014 to January 2015, which was available by April 2015^1–7^. A relatively large degree of sequence diversity is observed in this data set (Fig. 1). However, we find no evidence of positive selection within the EBOV-Makona outbreak, with neither Slr nor PAML identifying a signal at statistical significance. Sequence variability is highest in intergenic regions as these regions presumably have lower evolutionary constraint than protein-coding regions. Within the protein-coding genes, disordered regions accommodate a disproportionate amount of EBOV’s nonsynonymous variation (Fig. 1). In these regions the general characteristic of the protein sequence is more important than residues at particular sites; a much wider range of sequences are therefore compatible, and purifying selection is relaxed resulting in higher evolutionary rates. This lowered constraint allows more replacements in these regions, a signal not to be confused with positive selection^5,16^. Importantly, lack of evidence for positive selection (average nonsynonymous to synonymous substitution ratios <1, Fig. 1), does not necessarily mean a lack of functional change. For example, residue replacements in intrinsically disordered regions of EBOV proteins GP and NP (Fig. 2 and 3) may affect both protein-protein interactions and the host immune response. The disordered mucin-like domain in GP constitutes approximately one third of the protein and so is a major constituent of the external surface of the virus, a primary target of immune surveillance. Disordered regions do not have a single unique structure, but nonetheless have been implicated in virulence^17^. In particular the conserved short linear motifs (SLiMs)^18^ form molecular interactions^19,20^. SLiMs are found within the disordered regions (Fig. S1) and may permit EBOV to form novel host interactions via “sticky” non-specific interactions with host proteins^21^, and so have the potential to randomly explore novel host perturbations.

**Fig. 1.**
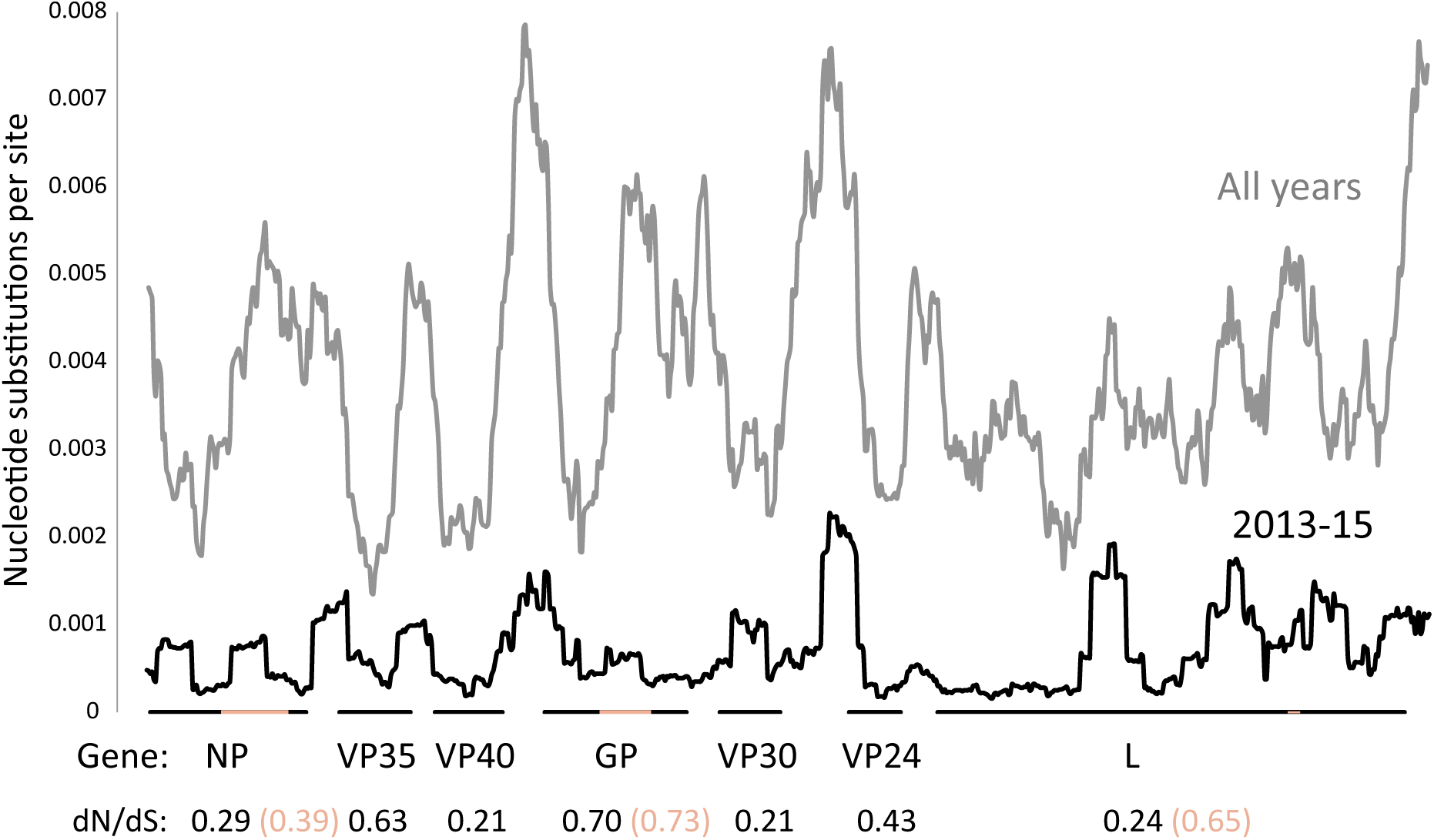
Genetic diversity (substitutions/site) plotted as a 500 nucleotide sliding window at 25 nucleotide increments, using a Jukes-Cantor model in DNASp (http://www.ub.edu/dnasp/), for 2013-2015 genomes only (black) and all genomes since 1976 (grey). Average rate of non-synonymous to synonymous substitution (dN/dS) is given for each gene (black) or shorter disordered regions (beige).

**Fig. 2.**
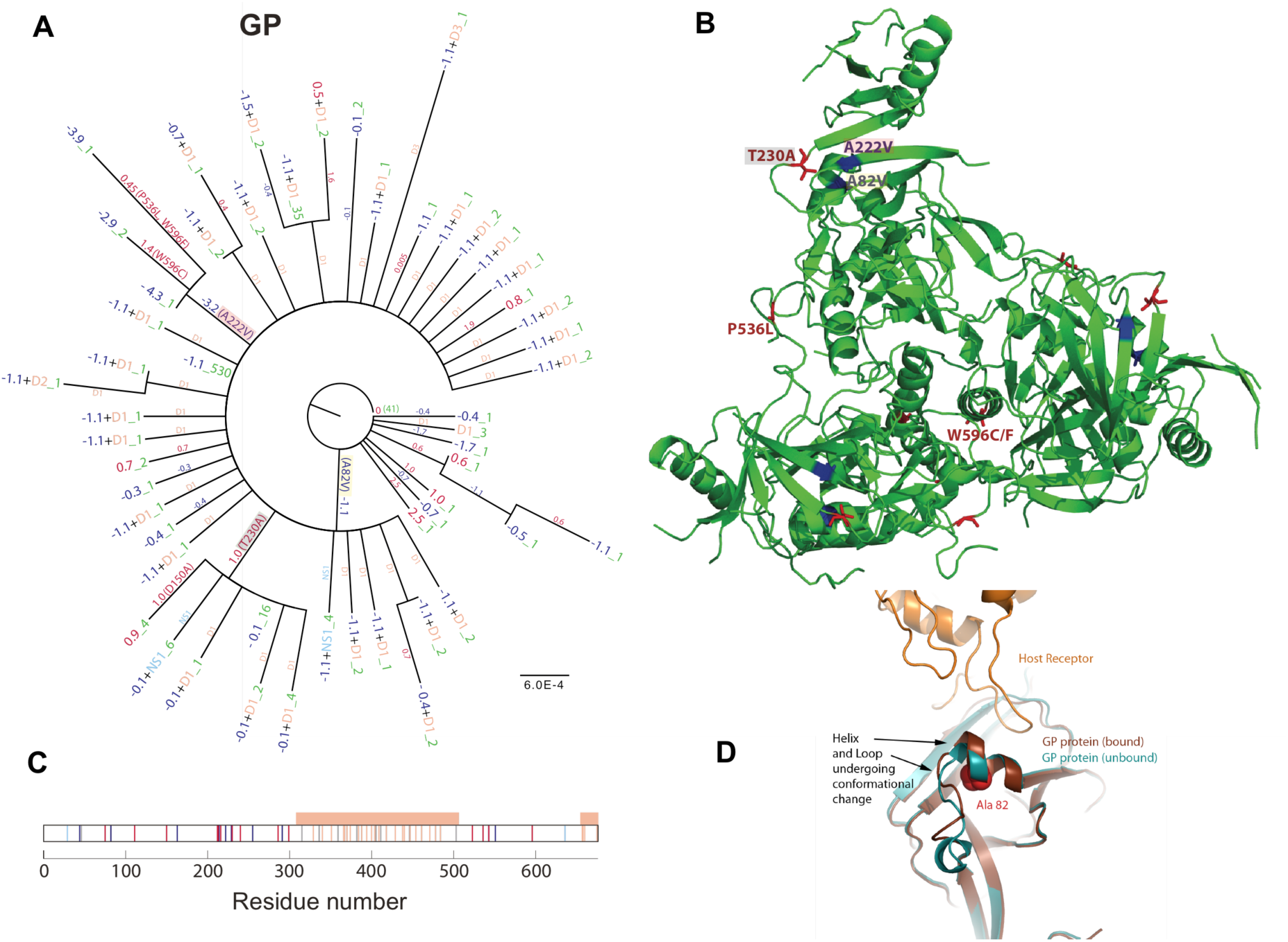
Phylogenetic trees inferred from amino acid sequences from the 2013-2015 EBOV outbreak (A) showing the impact of amino acid replacements on the stability of the GP protein structure. The likely ΔΔGs (in kcal/mol) for all cumulative replacements are shown at the tips, with positive numbers (destabilising) coloured red and negative numbers (stablising) in blue. Where a sequence is present more than once in the population, the number of occurrences is shown in green. Where replacements fall in disordered regions or ordered regions with no known structure (denoted by D (beige) or NS (pale blue), respectively), ΔΔG values cannot be estimated; the cumulative number is listed after the letter D or NS. The scale bar indicates the number of amino acid replacements per site. The location and type of selected residue changes (see Results for further details) are highlighted on the NP protein structure (B); red and blue colours indicate destabilising and stabilising, respectively. The location of the residue changes are shown (C), where red and blue bars indicates destabilising and stabilising, respectively; beige bars indicate disordered regions (indicated by beige rectangle). The proximity of the Ala 82 to Val replacement and the host receptor and the conformational change on receptor binding is also shown (D).

**Fig. 3.**
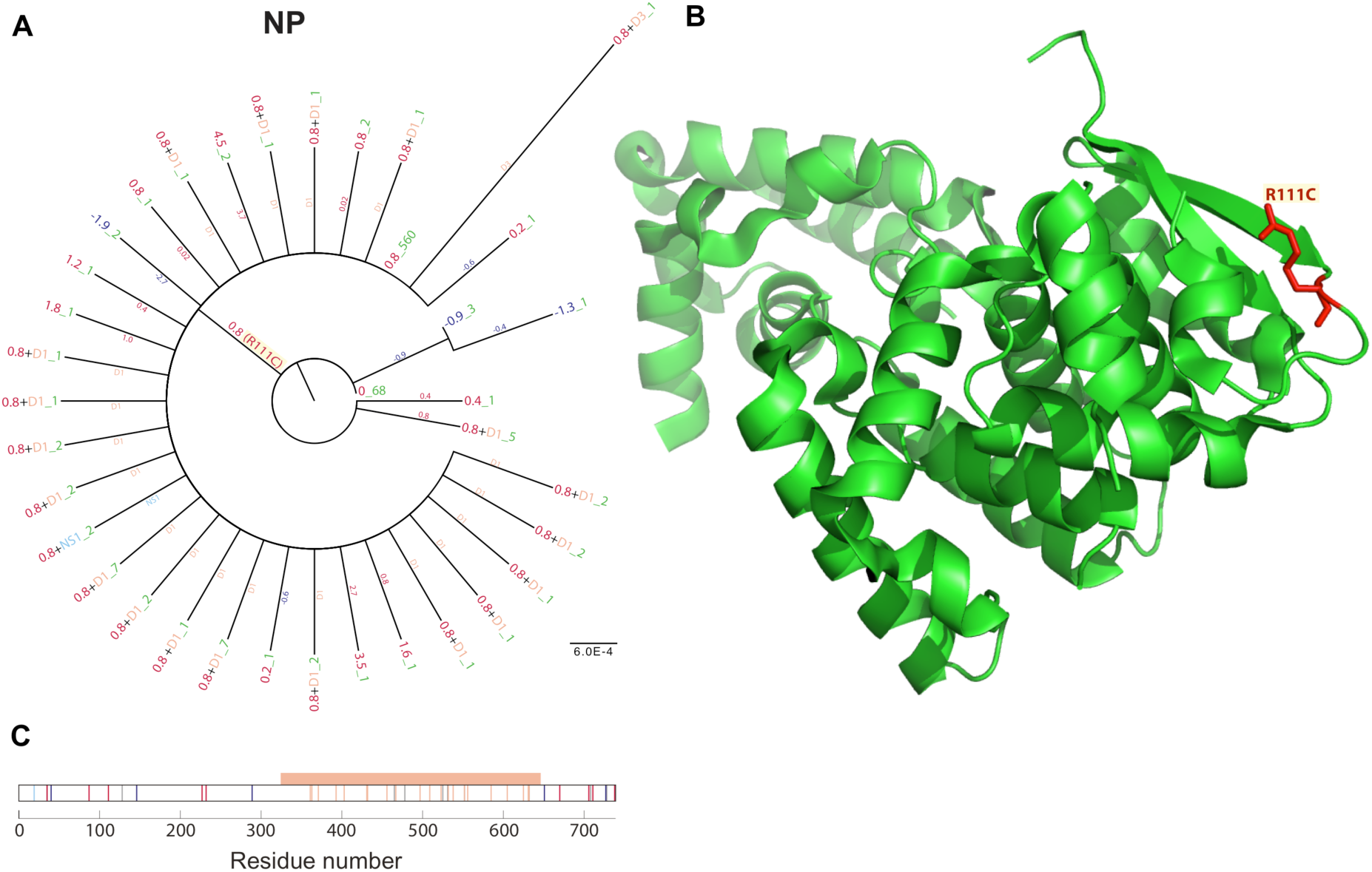
Phylogenetic trees inferred from amino acid sequences from the 2013-2015 EBOV outbreak (A) showing the impact of amino acid replacements on the stability of the NP protein structure. See figure 2’s legend for further details. The location and type of selected residue changes are highlighted on the GP protein structure (**B)**. The location of the residue changes are shown (C), where red and blue bars indicates destabilising and stabilising, respectively; beige bars indicate disordered regions (indicated by beige rectangle).

To analyse and compare variation in EBOV proteins, molecular phylogenetic trees were constructed and ancestral genetic sequence changes reconstructed. By modelling EBOV sequences in the context of their three-dimensional proteins we studied the nature of the amino acid residue replacements (non-synonymous substitutions) through time on the phylogenetic tree. Replacements tend to be sparsely dispersed across the variants sampled, explaining the lack of structure in the inferred amino acid trees for all proteins (Fig. 2-4), with only a minority of changes being present in transmitted viruses.

**Fig. 4.**
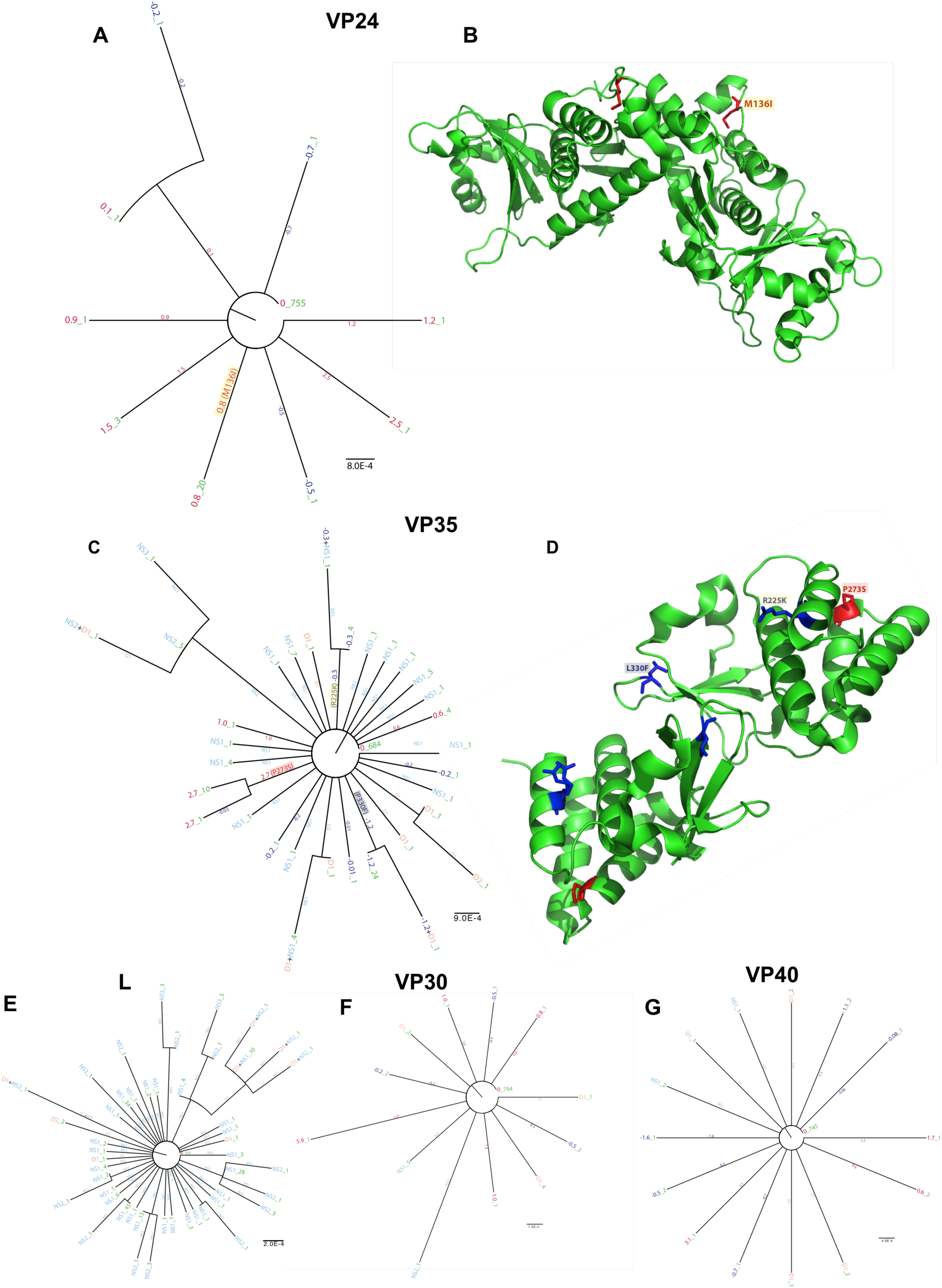
Phylogenetic trees inferred from amino acid sequences from the 2013-2015 EBOV outbreak showing the impact of amino acid replacements on the stability of the VP24, VP35, L, VP30 and VP40 protein structure (panels A, C, E, F and G, respectively). See figure 2’s legend for further details. The location and type of selected residue changes occurring in the VP24 and VP35 protein structure (panels B and D, respectively).

In the GP protein we find 6 variants (Fig 2A) in the structured region. Two of these (replacement of Ala with Val at residue 222, and replacement of Thr 230 with Ala highlighted in red and grey respectively in Fig. 2A and B) are in the glycan cap. The replacement at position 222 is predicted to have a relatively strong stabilising effect with a ΔΔG of -3.2 kcal/mol, whereas that at position 230 is mildly destabilising. The glycan cap, along with the mucin domain, is cleaved to activate the GP protein^22^. Replacement of Ala with Val at residue 82 (highlighted in yellow in Fig. 2A) appears to be the most significant replacement. This variant (KR817182 – patient EM_0794104) was first detected in Gueckedou Guinea on 31^st^ March 2014, before the spread to Sierra Leone, and corresponds to a nucleotide substitution of C to T at site 6281 (Fig. S2) and is in the last common ancestor leading to the major Lineage B of the outbreak^4^, consistent with a selective sweep (Fig. 2A). Previous analysis of amino acid replacements did highlight this change^9,23^, and Ladner *et al.*^8^ identified it being subject to positive selection, and its proximity to the receptor binding domain. Subsequent determination of the crystal structures of the host-primed GP protein in both the uncomplexed form ^24^ and bound to the host receptor NPC1 ^25^ allow additional analysis.

The priming of the GP protein involves cleavage of the glycan cap, which reveals a hydrophobic binding pocket ^24^. Comparison of the unprimed, primed/unbound and primed/bound structures (Fig. 2D) (PDB codes 3CSY, 5HJ3 and 5F1B) shows a conformational change ^25^, which occurs upon binding to the receptor, rather than upon cleavage of the glycan cap. The residues undergoing the binding-specific conformational change are a short loop (residues 71-78) and the subsequent 3_10_ helix (79-86). The A82V replacement does not make direct contact with the host receptor, but rather is in an adjacent position within the helix that changes conformation relative to the rest of the GP protein. A range of variants within this region have been demonstrated to alter binding affinity between the virus and receptor ^26^. This replacement (Fig. 2B) is predicted to stabilize the protein in all three available structures (ΔΔG of -1.1, -1.6 and -0.8 kcal/mol in the unprimed, primed/unbound and primed/bound respectively), probably through the additional van der Waals interactions formed with neighboring residues. The comparison of the ΔΔGs for the bound and unbound states allows a prediction of the effect of the replacement on binding. Both forms are stabilised relative to the wild type, but the unbound state more so. This indicates the A82V replacement shifts the equilibrium in the direction of the unbound form, decreasing the binding affinity of the virus for the human host receptor.

We also observe a small number of strains in which GP is predicted to have increased thermal stability (detailed Fig. 2A). These strains are frequently characterised by the combination of stabilising changes that precede later destabilising changes. This pattern of replacements demonstrates how stabilising replacements are potentially important as they can permit the accommodation of otherwise deleterious destabilising replacements, as has been demonstrated in enabling drug or antibiotic resistance to evolve^10,11,27,28^ Indeed in the GP phylogeny there is a cluster of six variants (corresponding to 33 infections) characterised by a destabilising replacement (replacement of Thr with Ala at residue 230, highlighted in grey in Fig. 2A and B) with one descendent accommodating a further destabilising change (replacement of Asp with Ala at residue 150). The Thr to Ala substitution was first detected in Moyamba, Sierra Leone on 4^th^ August 2014 (KR105267, patient G4725.1) and subsequently spread to Guinea, first detected there on 20^th^ August 2014 (KR534517, patient Conakry-684). The T230A clade was last detected in Sierra Leone near the capital Freetown on 11^th^ November 2014 (KP759705, patient Makona-J0168^3^) and in Forecariah in Guinea on 20^th^ October 2014 (KR534566/Guinea/2014/10/20, patient Makona-Forecariah-1571^29^) The further destabilising change within this clade, Asp with Ala at residue 150, was first detected in Port Loko, Sierra Leone on 30^th^ September 2014 (KP759691, patient Makona-J0032^3^) and last detected in patient Makona-J0168 (as above).

For NP the majority of variants exhibit destablising replacements (Fig. 3A) in the structured regions (positive ΔΔGs). The most significant replacement is Arg with Cys at residue 111 (highlighted in yellow in Fig. 3A) with a predicted positive free energy of folding (ΔΔG of 0.8 kcal/mol). This replacement, similar to A82V, arose early in the major B lineage of the outbreak, first detected in three patients sampled on 26^th^ May 2014 at Kissi Teng (Kailahun) in Sierra Leone (KM034551, KM034552 and KM034556; patients EM096, EM098 and G3677.1^1^). It corresponds to a nucleotide change from C to T at site 798In the context of the three-dimensional structure of the NP protein (Fig. 3B) this residue is on the surface of the molecule, and both the ancestral residue and the replacement make few, if any, intra-molecular interactions; this substitution is, thus, probably an unconstrained “neutral” substitution. Each individual NP sequence tends to display only one or two destabilizing mutations and in general sequences have moderately increased stability compared with ancestors (cumulative ΔΔGs in the range of +0.02-4.5 kcal/mol, compared with typical protein folding energies of 15-30 kcal/mol).

In VP30 and VP40 there is a clear predominant variant with no clusters of more than two variants identifiable (Fig. 4F and G). Interestingly in VP24 there is a destabilising replacement (Met with Ile at residue 136, highlighted in yellow Fig. 4A), which reduces the overall ΔΔG to 0.8 kcal/mol that corresponds to a cluster of 20 infections.

This slightly destabilising replacement is near to the protein surface (Fig. 4B). As such, these residues are likely to be functionally equivalent. In VP35 there are residue changes that cluster in the phylogenetic tree, indicating that they might be functionally important. These replacements correspond to both stabilising (Arg with Lys at residue 225 and Leu with Phe at residue 330, highlighted in yellow and grey respectively, Fig. 4C) and destabilising (Pro with Ser at residue 273, highlighted in red Fig. 4C) changes that correspond to 5, 25 and 11 infections, respectively. These replacements are dispersed in the protein structure, although all three are close to the protein’s surface (Fig. 4D). Of note this stabilising change in VP35 (corresponding to 25 infections) is embedded in the same cluster of viruses in VP24 characterised by a destabilising replacement (corresponding to 20 infections). Finally, in the L protein there are some sub-clusters (Fig. 4E), however, no structures are available to look at their context of these residue changes.

## Discussion

Throughout the EBOV-Makona epidemic there was extensive monitoring for altered viral properties that could explain EBOV’s “success” in comparison to previous outbreaks. None was found. In particular there was much focus on shifts in mutation rates and positive selection as an indicator of important change in the virus population. This emphasis is potentially misleading, since adaptation can arise by chance in a population regardless of the rate of evolution. Additionally concern had been raised about EBOV becoming adapted to humans via unspecified gain-of-function mechanisms, failing to recognize that a directional evolution can be associated with rare mutations that can be hard to detect in the case of a selective sweep, or that a less virulent, attenuated, virus would be at an advantage in the human population Previous detection of positive selection^30^ in EBOV-Makona, used FUBAR/HyPhy, whereas we use the more conservative methods implemented in Slr and PAML. The lack of statistically significant positive signal in our analysis demonstrates that the evidence for positive selection on the virus during the course of the EBOV-Makona outbreak, is borderline and dependent on the method of calculation used.

Our analysis of possible consequences of residue changes on protein structure confirms that EBOV is functionally very similar across the EBOV-Makona outbreak, with the observed nonsynonymous substitutions being mostly slightly deleterious in nature. Other amino acid replacements show a pattern of stabilising replacements preceding destabilising ones. EBOV amino acid replacements that have a deleterious effect on protein folding, for example as we have identified in GP and VP35 (Fig. 2 and 4), although we might expect them to be removed through purifying selection^1,5^, may not in fact reduce viral fitness in a human infection. Nonsynonymous variation was, however, accumulating in EBOV’s genome, albeit mainly in the intrinsically disordered regions of proteins. This cryptic variation has the potential to contribute to EBOV’s evolutionary capacitance as any novel variant may be better adapted to human transmission, and this could be the result of either a gain-of-function or an attenuated virus.

Alterations in receptor binding affinity arising from point mutations can influence the outcome of infection ^26^. Here, new crystal structures ^24,25^ have allowed us to identify sequence variants close to the host-receptor binding site. The comparison of the GP protein in the unprimed state ^22^ with both the primed ^24^ and primed and bound states ^25^ (Fig. 2D) allow not only the identification of a conformational change in the GP protein, but also allow this change to be associated with binding to the host receptor rather than with cleavage of the cap domain. Several point mutations and naturally-occurring replacements around the binding site have been shown to alter binding properties. These include variants in both the receptor and GP protein ^26^, which can both increase or decrease binding affinity. For example mutation of Thr 83 to Met (i.e. the site neighbouring the replacement we observe Ala 82 to Val) reduces binding, whereas mutation of Ala 141 to Val increases it ^26^. The replacement of Asp 502 to Phe in the host receptor NPC1 is observed in the African Straw-coloured fruit bat, and protects this species from EBOV. Inspection of the crystal structure of the complex suggests that this replacement removes a charge-charge interaction between Asp 502 in the receptor and Lys 155 in the GP protein, replacing it with the weaker van der Waals interaction. The GP Ala 82 to Val replacement occurred in the GUI-1 lineage found in Guinea. sampled in early 2014^31^. The earliest wave of Liberian cases also came from within this lineage^32^. This variant is one of only two amino acid replacements to have arisen in the latest epidemic and to have become fixed (Fig. S2) ^23^. It is located in the 310 helix that undergoes conformational change upon binding, and is predicted to stabilise the unbound state. If this prediction is correct, this change will result in a decreased affinity of the virus for the host receptor NPC1.

EBOV infection is usually sufficiently severe that patients are immobilized and can be readily identified. If a less virulent variant were to “seed” a cluster of infections with a higher reproductive number (R) due to it being less immobilising and/or identifiable in an infected person the result would be a wider and more rapid spread through the human population. Indeed there is evidence that EBOV is both less virulent in comparison to a previous outbreak^13^ and has decreased in virulence as the EBOV-Makona epidemic progressed^15^. There is also evidence for persistence infection of EBOV survivors for up to a year and a half, including transmission in this time^33.^

The risk of viral attenuation, and corresponding adaptation of EBOV to humans, necessitates both monitoring and more research. Results from our analysis are likely to be conservative because much of the available data was sampled from the sickest individuals. Typically, virus is characterised in patients with high levels of viral RNA as assessed with a real-time quantitative reverse transcriptase polymerase chain reaction (qRT-PCR)–based threshold cycle (CT) value^34^. For example, in one of our main data sets^4^ infected patients were chosen with a CT value of 20 or below and for these patients the survival rate was 20%.

Interestingly, bats, the probable host reservoir of EBOV, have a peak body temperature of ~42°C due to the metabolic requirements of flight^35^. In human infections, EBOV is thus replicating at lower temperatures than its usual bat host. This new niche for EBOV has potentially important consequences for protein folding/stability, the tolerance for change and ultimately viral fitness in the new host species. If transmission to humans is relaxing structural/functional constraints on EBOV proteins, e.g., for thermal stability, less virulent variants could emerge associated with deleterious change. These variants could be at least partially controlled by fever, and so become less debilitating.

In conclusion, our results demonstrate that EBOV was relatively stable in the human population. Coupled with the lack of evidence for change across previous outbreaks^36^ this implies EBOV, unlike most viral cross-species transmissions, does not require significant adaptation^37^ to replicate efficiently in humans. On this occasion, EBOV was prevented from establishing in the human population and becoming endemic, by drastic intervention strategies and changes in human behaviour that were feasible due to the relative ease of identifying infected individuals without any requirement for testing. It cannot be assumed that high virulence will always be the case for future EBOV outbreaks. Continued monitoring for outbreaks and rapid intervention is, thus, an imperative if we are to prevent EBOV becoming endemic in humans.

## Online Methods

Viral genomes were obtained from blood taken from 179 patients in Guinea and sent to the European Mobile Laboratory (EMLab) for sequencing (see ref^4^ for full details). This data was combined with the available EBOV genome sequence data from the EBOV-Makona outbreak (476 genomes)^1–7^ and sequences containing stop codons within genes or sequence ambiguities discarded. The remainder were aligned using MAFFT v.7.215 (http://mafft.cbrc.jp/alignment/software) and non-coding regions removed using MEGA v.6 (http://www.megasoftware.net). The alignment was then partitioned into individual alignments for each gene and duplicate sequences removed. All alignments are available from DOI: 10.17635/lancaster/researchdata/6. Positive selection analysis was carried out using PAML v.4.7 (http://abacus.gene.ucl.ac.uk/software/paml.html) and Slr (http://www.ebi.ac.uk/goldman-srv/SLR/). Amino acid sequence phylogenies were inferred with the WAG substitution model using RAXML (http://sco.h-its.org/exelixis/web/software/raxml). Ancestral EBOV sequences were estimated using maximum likelihood with FastML (http://fastml.tau.ac.il). Trees are rooted on the earliest sequences as denoted by sampling date. Energy changes for observed amino acid replacements were predicted using an empirical force field with FoldX version 3 Beta 6 (http://foldx.crg.es). See Olabode *et al*. (2015) for further details^36^. IUPred was used to predict disorder (http://iupred.enzim.hu) and conserved short linear motifs (SLiMs) predictions were made with ANCHOR^18^ using the default probability cut off value 0.5.

## Acknowledgements

We gratefully thank Kristian Andersen for pointing out the significance of the A82V residue change. Thanks to Dave Thornton for discussion on mucins. DM, JAH, SG, and MWC are funded by the EVIDENT (Ebola virus disease: correlates of protection, determinants of outcome, and clinical management) project funded by European Union’s Horizon 2020 research and innovation programme under grant agreement no. 666100. AO received support from the Faculty of Life Sciences, University of Manchester. XJ was supported by MRC (G1001806/1) and Wellcome Trust (097820/Z/11/B) funding to DLR. DG is supported by an Early Career Small Grant from Lancaster University.

---

**Supplementary figure S1.**
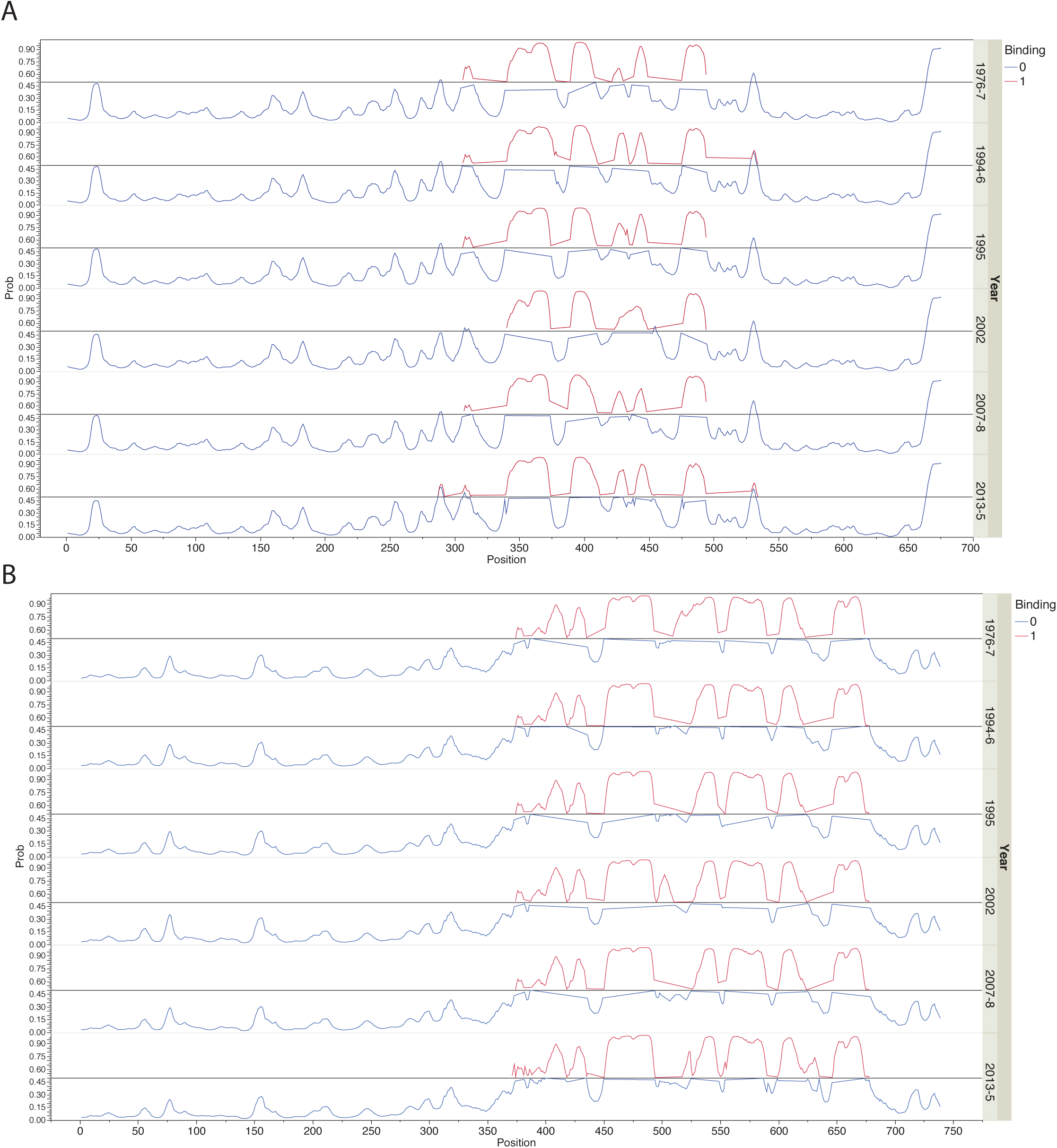
Prediction of binding regions (SLiMs) in intrinsically disordered regions in GP (A) and NP (B) for each EBOV outbreak. Red lines indicate the region is likely to bind to other proteins while blue lines indicates the region is not likely to bind. Straight lines link different red curves.

**Supplementary figure S2.**
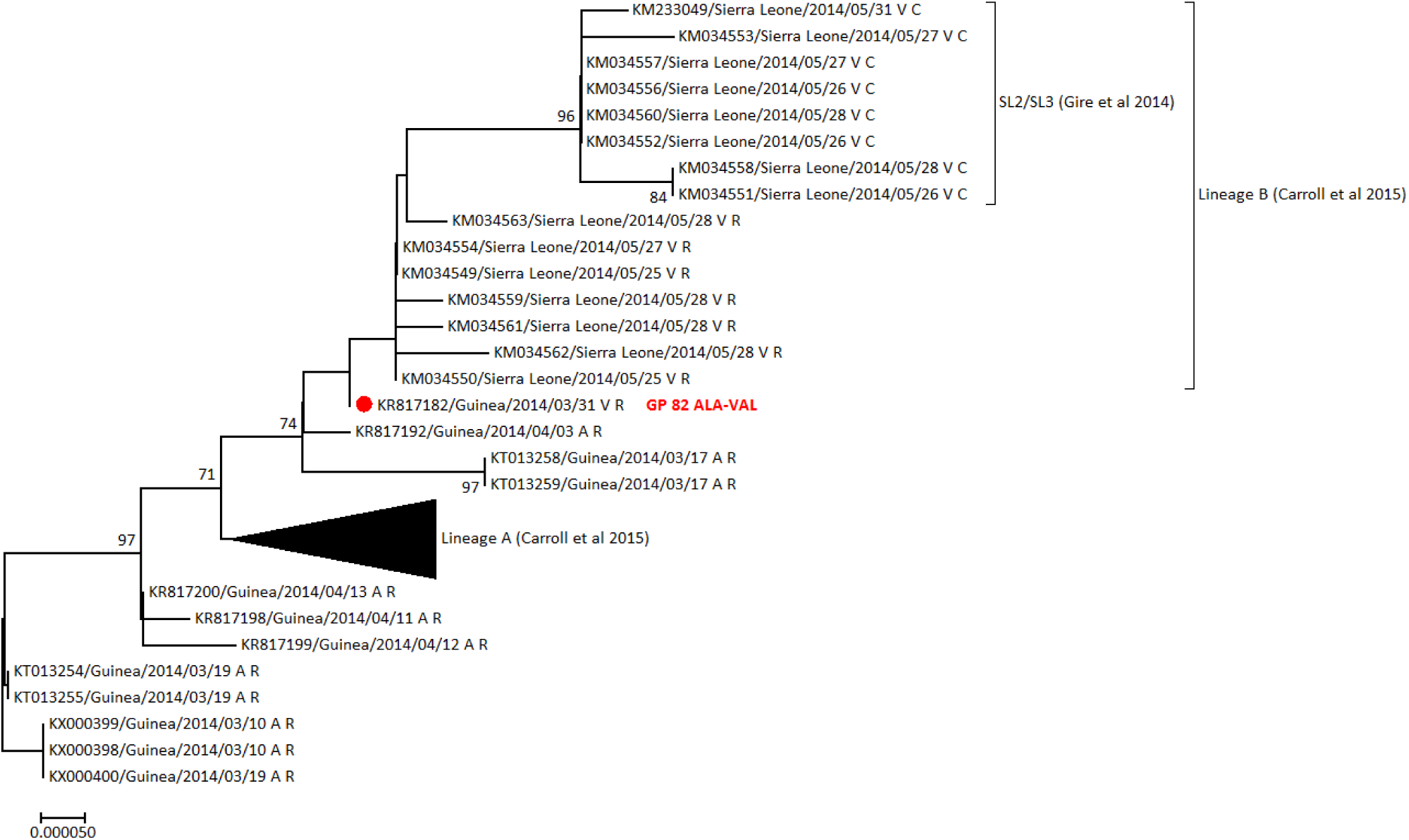
Maximum likelihood tree drawn in MEGA, with bootstrap confidence values indicated where >70, using EBOV-Makona full genome sequences collected from 10th March 2014 to 28th May 2014. Lineage A (Carroll et al 2015) which subsequently spread as a minor variant through Guinea, is collapsed. Lineage B (Carroll et al 2015) and its sublineage SL2/SL3 (Gire et al 2014) are indicated. Genomes are annotated with two letters referring to the amino acids at positions 82 of the glycoprotein and position 111 of the nucleoprotein respectively. Inital genomes had A at GP82 and R at NP111 and are therefore annotated "A R". Sequence KR817192 collected on 3rd April 2014 in Guinea (red dot) was the earliest with the GP82 A->V substititution "V R". The SL2/SL3 clade (Gire et al 2014) represent the subsequent NP111 R->C substitution "V C", occurring on or before 26th May 2014 in Sierra Leone.

## References

1. Gire, S.K., et al. Genomic surveillance elucidates Ebola virus origin and transmission during the 2014 outbreak. Science 345, 1369–1372 (2014).

2. Hoenen, T., et al. Complete genome sequences of three ebola virus isolates from the 2014 outbreak in west Africa. Genome announcements 2(2014).

3. Tong, Y.G., et al. Genetic diversity and evolutionary dynamics of Ebola virus in Sierra Leone. Nature (2015).

4. Carroll, M.W., et al. Temporal and spatial analysis of the 2014-2015 Ebola virus outbreak in West Africa. Nature (2015).

5. Park, D.J., et al. Ebola Virus Epidemiology, Transmission, and Evolution during Seven Months in Sierra Leone. Cell 161, 1516–1526 (2015).

6. Hoenen, T., et al. Virology. Mutation rate and genotype variation of Ebola virus from Mali case sequences. Science 348, 117–119 (2015).

7. Kugelman, J.R., et al. Monitoring of Ebola Virus Makona Evolution through Establishment of Advanced Genomic Capability in Liberia. Emerging infectious diseases 21, 1135–1143 (2015).

8. Ladner, J.T., et al. Evolution and spread of Ebola virus in Liberia, 2014-2015. Cell host & microbe 18, 659–669 (2015).

9. Olabode, A.S., Jiang, X., Robertson, D.L. & Lovell, S.C. Ebolavirus is evolving but not changing: No evidence for functional change in EBOV from 1976 to the 2014 outbreak. Virology 482, 202–207 (2015).

10. DePristo, M.A., Weinreich, D.M. & Hartl, D.L. Missense meanderings in sequence space: a biophysical view of protein evolution. Nature reviews. Genetics 6, 678–687 (2005).

11. Tokuriki, N. & Tawfik, D.S. Stability effects of mutations and protein evolvability. Curr Opin Struct Biol 19, 596–604 (2009).

12. Kerr, P.J., et al. Evolutionary history and attenuation of myxoma virus on two continents. PLoS pathogens 8, e1002950 (2012).

13. Marzi, A., et al. Delayed Disease Progression in Cynomolgus Macaques Infected with Ebola Virus Makona Strain Emerging infectious diseases 21, in press (2015).

14. Lanini, S., et al. Blood kinetics of Ebola virus in survivors and nonsurvivors. The Journal of clinical investigation 125, 4692–4698 (2015).

15. de La Vega, M.A., et al. Ebola viral load at diagnosis associates with patient outcome and outbreak evolution. The Journal of clinical investigation 125, 4421–4428 (2015).

16. Liu, S.Q., Deng, C.L., Yuan, Z.M., Rayner, S. & Zhang, B. Identifying the pattern of molecular evolution for Zaire ebolavirus in the 2014 outbreak in West Africa. Infection, genetics and evolution: journal of molecular epidemiology and evolutionary genetics in infectious diseases 32, 51–59 (2015).

17. Goh, G.K., Dunker, A.K. & Uversky, V.N. Detection of links between Ebola nucleocapsid and virulence using disorder analysis. Molecular bioSystems (2015).

18. Dosztanyi, Z., Meszaros, B. & Simon, I. ANCHOR: web server for predicting protein binding regions in disordered proteins. Bioinformatics 25, 2745–2746 (2009).

19. Meszaros, B., Simon, I. & Dosztanyi, Z. Prediction of protein binding regions in disordered proteins. PLoS computational biology 5, e1000376 (2009).

20. Davey, N.E., et al. Attributes of short linear motifs. Molecular bioSystems 8, 268–281 (2012).

21. Xue, B. & Uversky, V.N. Intrinsic disorder in proteins involved in the innate antiviral immunity: another flexible side of a molecular arms race. Journal of molecular biology 426, 1322–1350 (2014).

22. Lee, J.E., et al. Structure of the Ebola virus glycoprotein bound to an antibody from a human survivor. Nature 454, 177–182 (2008).

23. Giovanetti, M., et al. Amino acid mutations in Ebola virus glycoprotein of the 2014 epidemic. J Med Virol 87, 893–898 (2015).

24. Bornholdt, Z.A., et al. Host-Primed Ebola Virus GP Exposes a Hydrophobic NPC1 Receptor-Binding Pocket, Revealing a Target for Broadly Neutralizing Antibodies. MBio 7, e02154–02115 (2016).

25. Wang, H., et al. Ebola Viral Glycoprotein Bound to Its Endosomal Receptor Niemann-Pick C1. Cell 164, 258–268 (2016).

26. Ng, M., et al. Filovirus receptor NPC1 contributes to species-specific patterns of ebolavirus susceptibility in bats. Elife 4(2015).

27. Gong, L.I., Suchard, M.A. & Bloom, J.D. Stability-mediated epistasis constrains the evolution of an influenza protein. Elife 2, e00631 (2013).

28. Studer, R.A., Christin, P.A., Williams, M.A. & Orengo, C.A. Stability-activity tradeoffs constrain the adaptive evolution of RubisCO. Proc Natl Acad Sci U S A 111, 2223–2228 (2014).

29. Simon-Loriere, E., et al. Distinct lineages of Ebola virus in Guinea during the 2014 West African epidemic. Nature 524, 102–104 (2015).

30. Ladner, J.T., et al. Evolution and Spread of Ebola Virus in Liberia, 2014-2015. Cell host & microbe 18, 659–669 (2015).

31. Simon-Loriere, E., et al. Distinct lineages of Ebola virus in Guinea during the 2014 West African epidemic. Nature 524, 102–104 (2015).

32. Ladner, J.T., et al. Evolution and Spread of Ebola Virus in Liberia, 2014-2015. Cell Host & Microbe 18, 659–669 (2015).

33. Diallo, B., et al. Resurgence of Ebola virus disease in Guinea linked to a survivor with virus persistence in seminal fluid for more than 500 days. Clin Infect Dis (2016).

34. Grolla, A., et al. The use of a mobile laboratory unit in support of patient management and epidemiological surveillance during the 2005 Marburg Outbreak in Angola. PLoS neglected tropical diseases 5, e1183 (2011).

35. Brook, C.E. & Dobson, A.P. Bats as 'special' reservoirs for emerging zoonotic pathogens. Trends in microbiology 23, 172–180 (2015).

36. Olabode, A.S., Jiang, X., Robertson, D.L. & Lovell, S.C. Ebolavirus is evolving but not changing: No evidence for functional change in EBOV from 1976 to the 2014 outbreak. Virology 482, 202–207 (2015).

37. Parrish, C.R., et al. Cross-species virus transmission and the emergence of new epidemic diseases. Microbiology and molecular biology reviews: MMBR 72, 457–470 (2008).

